# State transitions in the statistically stable place cell population are determined by rate of perceptual change

**DOI:** 10.1101/2021.06.16.448638

**Authors:** Sander Tanni, William de Cothi, Caswell Barry

## Abstract

The hippocampus plays a central role in mammalian navigation and memory, yet an implementational understanding of the rules that govern the granularity of location encoding and the spatial-statistics of the population as a whole are lacking. We analysed large numbers of CA1 place fields recorded while rats foraged in environments up to 8.75 m^2^. We found that place cell propensities to form fields were proportional to open-field area, gamma-distributed, and conserved across environments. The properties of place fields varied positionally with a denser distribution of smaller fields near boundaries. Remarkably, field size and density were in a dynamic equilibrium, such that population-level activity statistics remained constant. We showed that the rate of transition through the statistically stable place cell population matched the rate of change in the animals’ visual scenes - demonstrating that the resolution of the spatial-memory system is bounded by perceptual information afforded by the environment.

## Introduction

Hippocampal place cells, pyramidal neurons in regions CA1 and CA3, are distinguished by their spatially constrained firing fields (O’Keefe and Dostrovsky, 1971). As a population, the activity of these cells provides a sparse representation of self-location that is relatively independent of other variables - such as head direction and velocity (Muller et al., 1994; Skaggs and McNaughton, 1998) - and is believed to provide the neural basis of a cognitive map (O’Keefe and Nadel, 1978). Fifty years of research have contributed extensively to our knowledge of this system and today place cells are understood to be common to mammals (Ekstrom et al., 2003; Yartsev and Ulanovsky, 2013), have been shown to be engaged as a temporal and abstract code (Grosmark and Buzsáki, 2016; Kraus et al., 2013), and are known to be reactivated during periods of quiescence (Ólafsdóttir et al., 2017; Wilson and McNaughton, 1993).

Despite these achievements, our understanding of the dynamics that control the statistics and distribution of place cell representations have been slower to advance. In part, this is due to technical barriers that make it difficult to collect long-duration, high-yield recordings in animals as they explore large spaces. Despite these constraints, a small number of groups have conducted work in extended environments, showing that individual place cells can develop multiple fields (Fenton et al., 2008) and exhibit distinct propensities to be recruited on long linear tracks (Lee et al., 2020; Rich et al., 2014). Similarly, investigations along the hippocampal axis identified a gradient of spatial scales, with ventral cells having considerably larger place fields than dorsal cells (Kjelstrup et al., 2008). Nevertheless, an implementational understanding of the hippocampal population code is still lacking (Marr, 1982) - effectively we know little about the rules that govern how the place code is recruited, how it evolves across space, and how it is influenced by sensory information.

One practical outcome of this situation is that we do not have sufficient empirical data to arbitrate between classes of computational models. For example, geometric cue-based models describe place field locations by integrating distance and direction from visual features such as environmental boundaries (Hartley et al., 2000; O’Keefe and Burgess, 1996). Since the rate of change of visual cues is proportional to their proximity, this suggests that the place code should vary more rapidly near to salient features. In contrast, models based on attractor dynamics tend to ignore any systematic variance in place field size and density across environments, emphasising even coverage and carefully balanced activity (Káli and Dayan, 2000; Samsonovich and McNaughton, 1997). These two classes of model, as well as others (de Cothi and Barry, 2020; Stachenfeld et al., 2017; Uria et al., 2020), provide competing but not incompatible explanations of hippocampal dynamics but the evidence needed to generate a synthesis is lacking.

Here we analyse large populations of place cells recorded while rats foraged in different-sized, equally-proportioned environments up to 8.75 m^2^. We find that place fields and cells are recruited in proportion to environmental area but with a strong influence of location on field frequency and size - fields are smaller and more numerous near boundaries, being larger and dispersed towards the environment centre. Surprisingly these two effects exactly counter each other, resulting in stable population-level firing within and between environments - suggesting the presence of a strong homeostatic mechanism governing place cell activity. Using a virtual reality (VR) replica of the recording environment, we show that the rate of change of the place cell population is strongly correlated with the rate of change in the animals’ visual scenes. Thus, while the summary statistics describing the distribution of place cell activity were stable across time and space, the rate at which this distribution evolved was dependent on the perceived granularity of the environment. Taken together these results suggest that the size and extent of individual place fields are well described by geometric cue-based models whereas the population as a whole conforms to the expectations of attractor-based models. More generally, as predicted by theory (Brown and Bäcker, 2006; Zhang and Sejnowski, 1999), it appears that the effective scale of representations within the spatial memory system are limited by the perceptible information afforded by the environment.

## Results

Using extracellular electrodes (128 channels per animal) we recorded 629 CA1 place cells (89-172 cells/rat) from five rats while they foraged for randomly dispensed reward (20 mg pellets) in four familiar, differently-sized environments. The environments - designated A to D - were identically proportioned with each being double the area of the previous to a maximum of 8.75 m^2^ (Fig. 1A, Fig. S1). The order of environments B, C and D was randomised for each animal, with the smallest environment A being used at the beginning and end of the recording session - recording duration was scaled proportional to environment size and a single recording session consisting of 5 trials was analysed from each animal (Fig. 1B). Place cells were isolated based on their waveforms (Fig. 1C) and temporal firing rate statistics (Fig. 1D) with many cells being active in multiple environments (Fig. 1E). Individual place fields were detected iteratively as contiguous regions of stable firing rates continuously increasing towards a peak (see Methods) (Fig. S2).

**Fig. 1.**
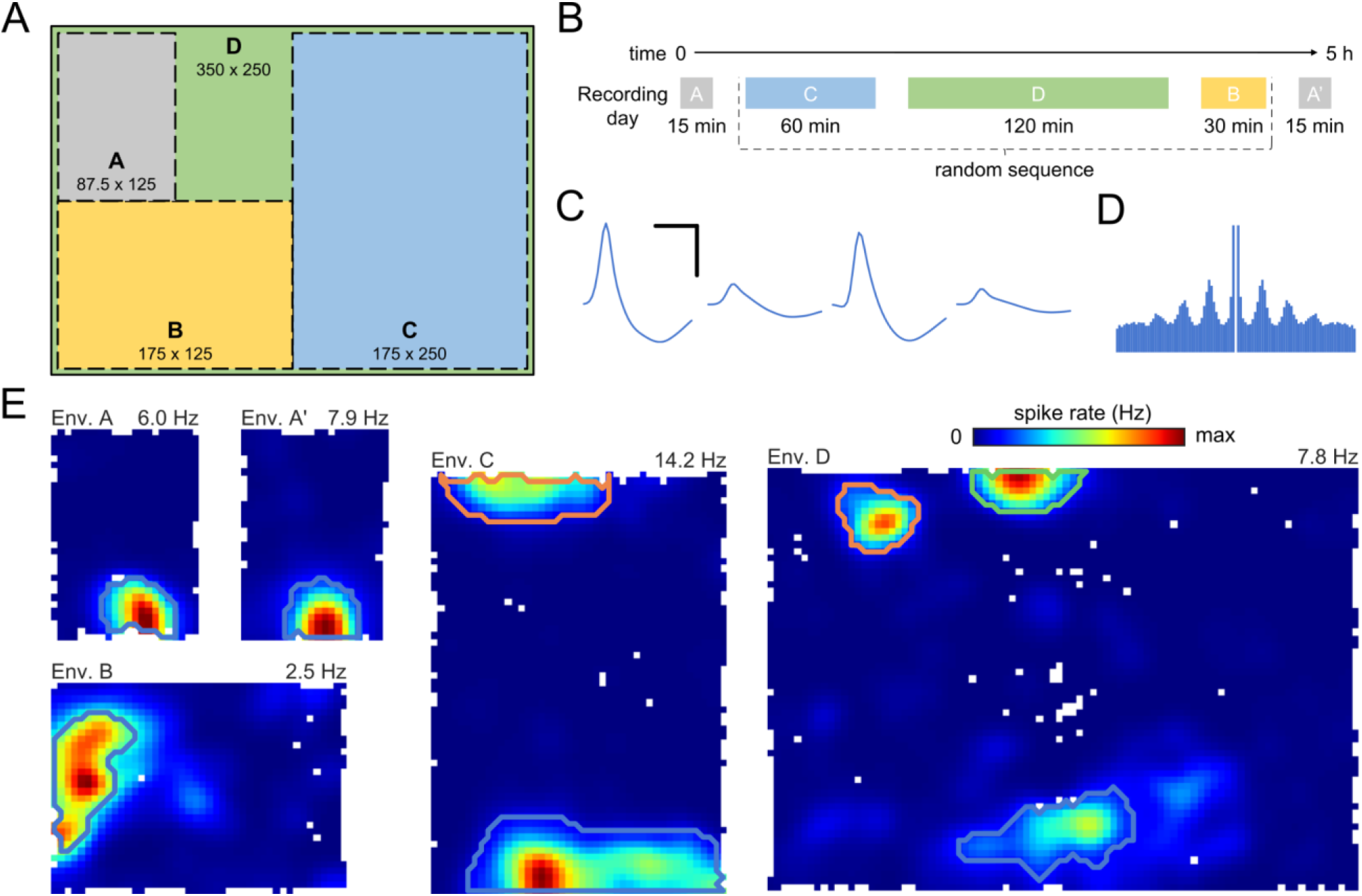
Place cell recordings in multiple large environments. (**A**) Schematic of the four environments (A, B, C and D) illustrating their relative sizes (cm) and positions in the experimental room. Each environment was distinguished by a set of unique cues. (**B**) Rats foraged in all environments during the recording session - environment A twice, at the start and end of the session, interleaved with the other three in random order. Recording duration scaled linearly with environment area. (**C, D, & E**) Waveforms (scale bars, 0.5 ms and 100 µV), autocorrelogram (maximum lag 500ms), and ratemaps for a typical place cell with activity in all environments (bin size 4 cm). Distinct place fields are delineated with lines of different colour. The colour map for each plot scales from 0 Hz to the peak rate above each map. Unvisited bins are white.

As is typical of the hippocampal system, the spatially modulated activity of neurons in environment A was stable both within and between recordings in environment A (Fig. S3A, intra-trial (A ½ v ½) spatial correlation: 0.61; inter-trial (A v A’): 0.51; inter-trial with shuffle (A v shuffled A’): -0.03) while activity in the four different environments was highly distinct, being sufficient to assign 94.5% of activity vectors (1s duration) to the correct environment (see Methods) (Fig. S3B).

In all animals, the number of active place cells - neurons with at least one spatial field - was greater in larger environments (Fig. 2A&E), as was the total number of place fields (Fig. 2B). However, place field recruitment increased more rapidly such that the average number of fields per cell was higher in larger environments (1.28 fields/cell in A to 2.28 fields/cell in D, Fig. 2C). Indeed, when all recording environments were considered collectively, representing a total space of 16.4 m^2^, it was more common for cells to have 2 fields than 1 and 84.8% of all cells had multiple fields (Fig. 2C). Furthermore, consistent with work conducted in 1D VR (Lee et al., 2020), we found that individual cells had different propensities to form place fields, and that this individual proclivity was maintained across environments (Fig. S4), meaning that cells with numerous fields in one environment were more likely to have numerous fields in another. Thus, the number of fields per cells were better fit with a gamma-Poisson model than equal-Poisson model (Rich et al., 2014) (log-likelihoods: -1538 vs -1679; Bayesian information criterion: 3089 vs 3365), with the latter generally overestimating the proportion of cells with 4-6 fields across the collective environment (Fig. S5). In turn, the same model accurately predicted the rate at which place cells were recruited (≥1 field) as a function of environment area (Fig. 2E), indicating that 99% of CA1 place cells will have at least one place field in environments greater than 51.8 m^2^ (Fig. 2E inset).

**Fig. 2.**
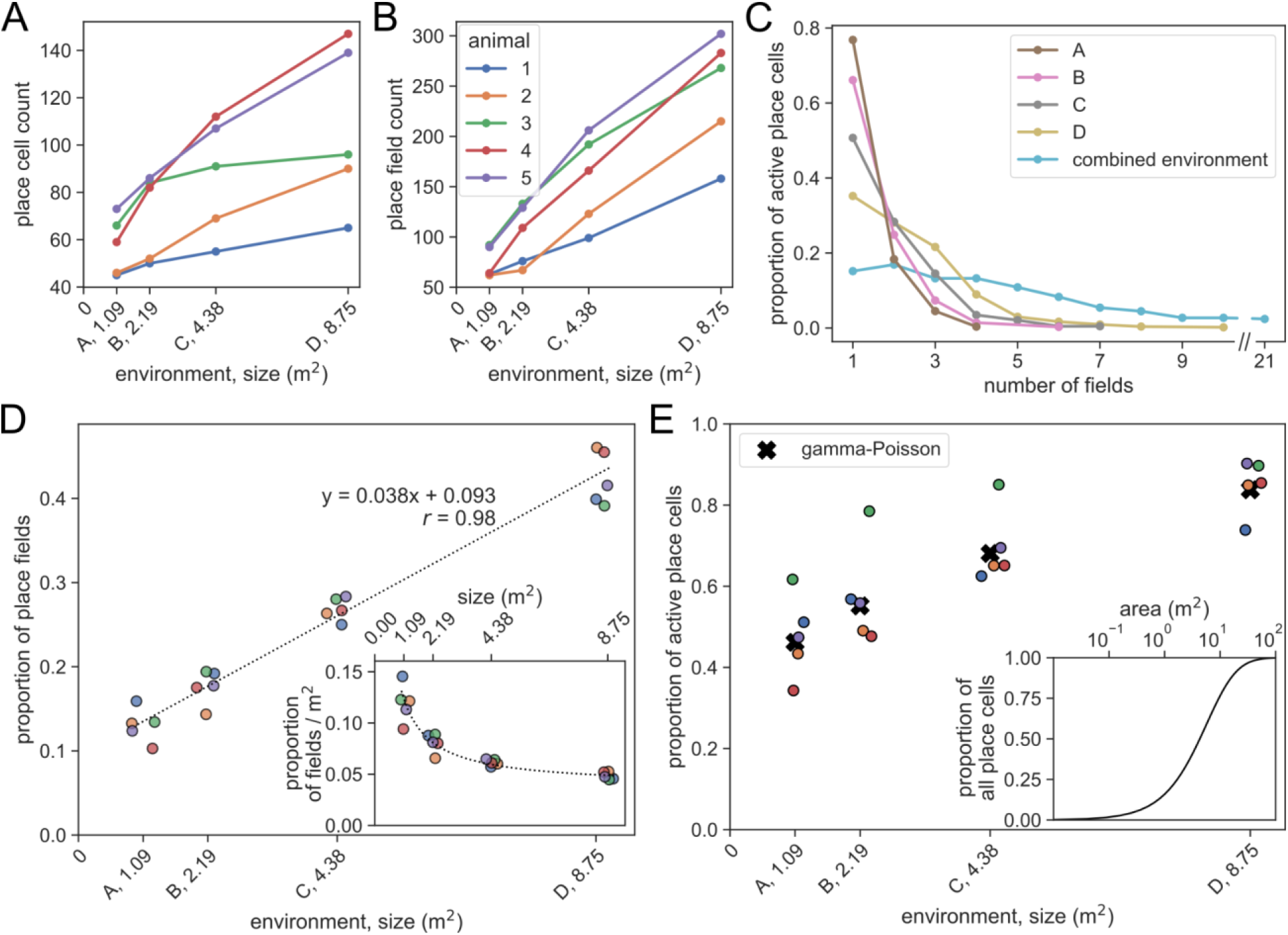
Place cells are more likely to be active and have more fields in large environments. Number of active place cells **(A)** and place fields **(B)** by environment area. (**C**) Distribution of place field counts per cell by environment. Only cells with at least one field in any environment are included. (**D**) Larger environments had more place fields, here shown as a proportion of all fields detected in each animal across the four environments (Kruskal-Wallis: H = 17.6, p = 5.36 x 10^-4^, N = 5), scaling linearly with environment area (dashed line, linear regression fit, *r* = 0.982, p = 10^-14^, N = 20). Inset shows place field count by environment area, regression line shown with same parameters as main plot (Kruskal-Wallis: H = 17.9, p = 4.71 x 10^-4^, N = 5). Animal colors same as **B**, points jittered to facilitate visualisation. (**E**) The proportion of all recorded place cells that were recruited in a given environment (≥ 1 field) increased with area (Kruskal-Wallis: H = 14.1, p = 0.003, N = 5) and was closely matched by a gamma-Poisson model fit to field numbers in a combined environment (see Fig. S5) adjusted by the relative field densities from Fig. 2D inset (see Methods) (mean squared error 0.0099). Inset shows the same gamma-Poisson model extrapolated to predict CA1 place cell recruitment in very large environments.

To quantify the relationship between the number of place fields and environment size, we examined how field recruitment varied with area, finding a remarkably robust linear regression with a positive intercept (*r* = 0.983, p = 10^-14^, slope = 0.038, intercept = 0.093, N = 20, Fig. 2D). Notably, this linear fit indicates that the proportion of place fields per unit area is lower in larger spaces (Fig. 2D inset, Kruskal-Wallis: H = 17.9, p = 4.71 x 10^-4^, N = 5).

Next, to understand why place fields were less numerous per unit area in large environments, we examined how fields were distributed within environments. To this end we segmented the space into concentric bands according to distance to the nearest boundary (each band 25 cm wide). In environments B to D, for which at least two bands could be defined, we calculated the density of place field peaks and found it was greater near to the walls - reducing towards the environment centre (Kruskal-Wallis tests: env. B, H = 4.81, p = 0.028; env. C, H = 12.5, p = 0.002; env. D, H = 16.1, p = 0.001; N = 5, Fig. 3A). Equally, considering only the band closest to the wall (<25cm), field peak density was generally higher in smaller environments (Fig. 3A, Kruskal-Wallis: H = 12.4, p = 6.02 x 10^-3^, N = 5). These effects could not be explained by the difference in dwell time between locations (Fig. S6). In direct contrast, the average size of place fields increased with distance from the wall (Kruskal-Wallis tests: env. B, H = 6.82, p = 0.009; env. C, H = 10.2, p = 0.006; env. D, H = 10.5, p = 0.015; N = 5, Fig. 3B) and were smaller in environment A (Kruskal-Wallis: H = 15.9, p = 0.001, N = 5, Fig. 3B). In particular, it appeared that the predominant factor contributing to this effect was that field width in a given axis was proportional to nearest wall-distance along that axis (Fig. S7) and not to wall-distance orthogonal to that axis (Fig. 3C&D, Kruskal-Wallis: H = 12.1, p = 0.007, N = 5; H = 0.39, p = 0.94, N = 5, respectively). So, compared to locations near the walls of an enclosure, there were on average fewer individual place fields further from the walls but those fields that were present tended to be larger.

**Fig. 3.**
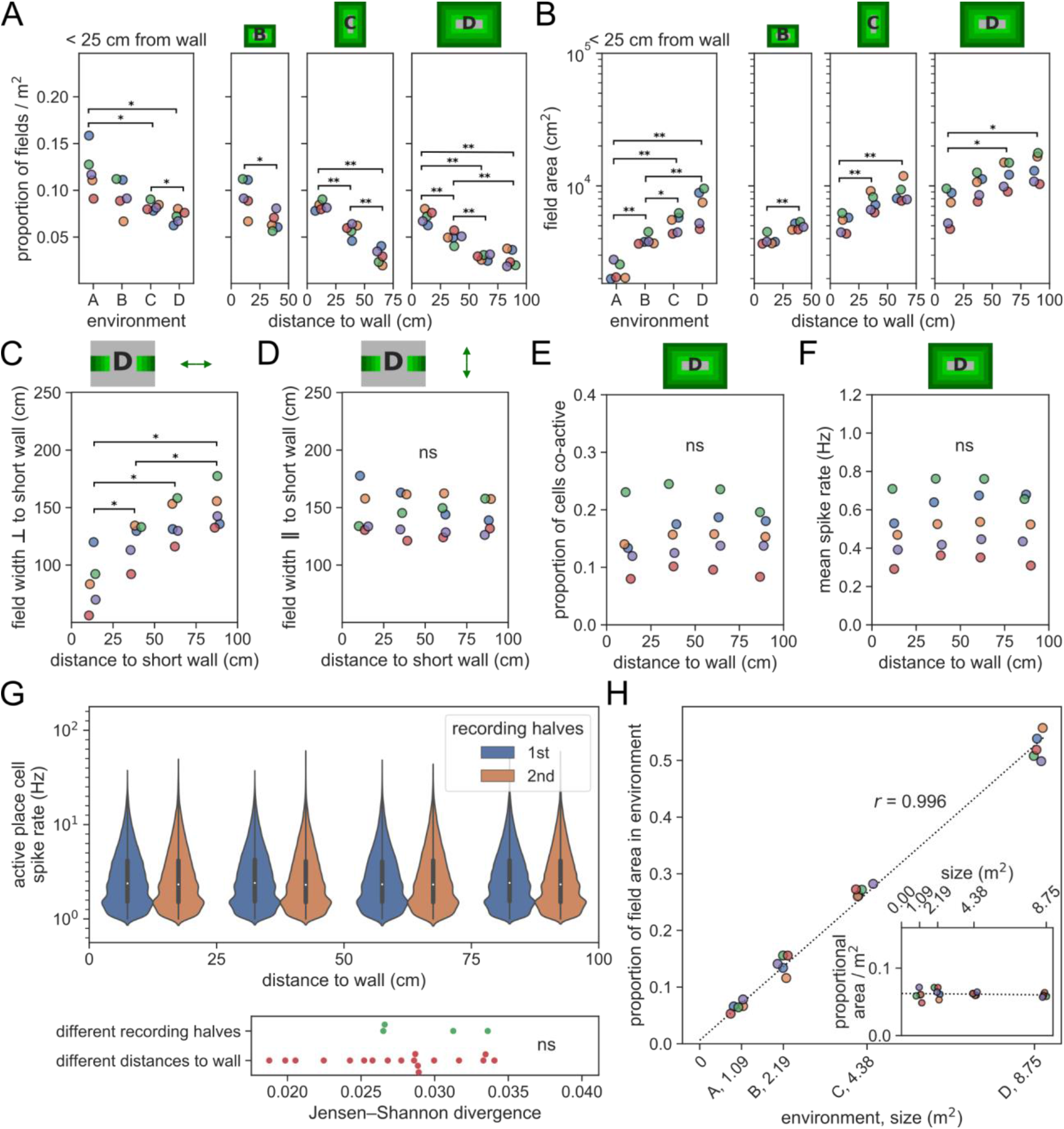
Density of place field peaks and the size of fields change with distance from the environmental boundary in a homeostatic manner. (**A**) Place field peak density per unit area (field peak/m^2^) is lower near the wall in large environments compared to small environments (left), and decreased with distance from the wall (right). Cartoon above each plot indicates wall distances in shades of green. Pairwise post hoc tests for this and subsequent panels adjusted for multiple comparisons using Benjamini/Hochberg (non-negative) correction. (**B**) The average area of place fields increased with distance from the wall and were larger in bigger environments when wall-distance was controlled for. (**C**) The average width of place fields measured orthogonal to the nearest wall (Fig. S7) in environment D increased with distance from the wall. (**D**) The average width of place fields measured parallel to the short wall was the same in all distance bins. The same observation was made in recordings from all environments (Fig. S8). (**E**) The proportion of active place cells (>1 Hz) was constant at different distances to the wall in environment D as was the mean firing rate of all place cells (**F**). (**G**) The distributions of firing rates at different distances to the wall in environment D (top) were stable across time and across distance bins (bottom). (**H**) The proportion of total place field area accounted for by place fields in each environment, computed separately for each animal, is highly correlated with the size of the environment (dashed line). Inset shows the values of the main plot divided by the size of each environment – the proportion of total place field area per m^2^ of each environment.

Considered alone, the observed decrease in the density of field peaks away from the boundary would be expected to result in fewer active place cells - yielding a lower mean firing rate. Conversely, the increase in field size would lead to greater overlap between place fields, resulting in the opposite outcome. Remarkably, we found these two effects were exactly balanced, meaning that neither the proportion of co-active place cells (firing rate > 1 Hz) nor the mean firing rate of the population varied with distance from the wall (Fig. 3E&F, Kruskal-Wallis: H = 1.2, p = 0.77 N = 5; H = 0.61, p = 0.89, N = 5; respectively). This was the case within and across all environments (Fig. S9). Similarly, interneuron firing rates were also constant at different distances to the bounding walls (Fig. S10, Kruskal-Wallis: H = 0.23, p = 0.97 N = 5). Furthermore, the distribution of firing rates in the place cell population was also unchanged at different distances to the wall (comparison of distribution divergence within and between locations Mann-Whitney: U = 45, p = 0.43, N = 4 and 18, Fig. 3G). These results are fundamentally underlined by a very high correlation between the environment size and the total place field area in that environment, expressed as a proportion of the total area of all fields recorded from an animal (linear regression: r = 0.996, p = 10^-20^, slope = 0.06, intercept = 0.006, N = 20) (Fig. 3H).

Thus, taken together, there appears to be several consistent and related features of the place cell code for space. First, the total area of place fields active in an environment is a near perfect linear function of the environment’s area. Second, population activity is homeostatically balanced, maintaining a constant proportion of active cells (15%) and mean firing rate (0.52 Hz/place cell) - despite field size growing with distance to walls. Finally, and more generally, the summary statistics describing the distribution of place cell activity is also stable across space.

Given that the distribution of firing rates in the place cell population is constant but the size of place fields and the density of their peaks varies systematically, this implies that hippocampal activity must evolve at different rates across the environment. Specifically, because place field width in a given axis is strongly determined by distance to the nearest wall orthogonal to that axis, the activity vector (see Methods on population activity change) should change fastest when the animal is near a wall and moving directly towards or away from it. Thus, analysing only trajectories running orthogonal to the short wall of the largest environment (D), we found the Euclidean distance between activity vectors (1 cm intervals) was greatest when animals were close to the wall (Kruskal-Wallis: H = 12.2, p = 0.007, N = 5) (Fig. 4A) an effect that was not found for parallel runs (Kruskal-Wallis: H = 3.78, p = 0.29, N = 5; difference between orthogonal and parallel, Mann-Whitney: U = 2, p = 0.018, N = 5, Fig. 4B). Similar effects were confirmed in the other environments (Fig. S11).

**Fig. 4.**
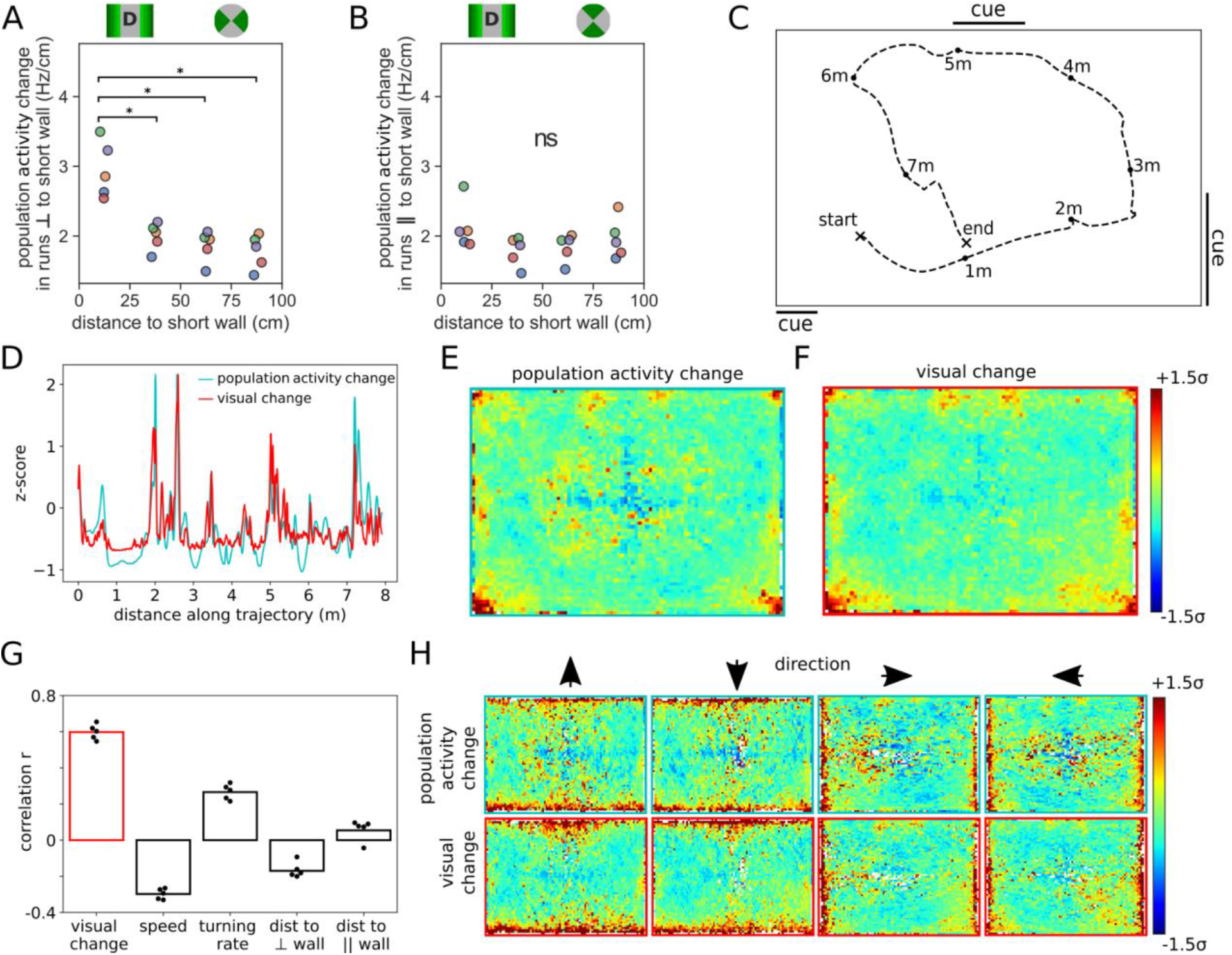
The rate of change in the place cell population mirrors the rate of change in the visual scene. Trajectories were split into 1 cm intervals and the Euclidean distance between adjacent activity vectors was calculated (see Methods on population activity change). (**A**) For trajectories orthogonal to the wall, the rate of change in the place cell population was greater when animals were close to the wall - an effect that is not observed for trajectories running parallel to the wall. Pairwise post hoc tests adjusted for multiple comparisons using Benjamini/Hochberg (non-negative) correction. (**B**). Legend above indicates spatial bins (left) and movement directions (right) used to plot data. (**C**) Example 8 m trajectory in the largest environment from animal 5. (**D**) Time series data for the trajectory in **C** showing a tight coupling between z-scored rate of change in the hippocampal population and reconstructed visual scene. (**E&F**) Spatially averaged rates of change for place cell population and visual scenes (data from all rats, no smoothing) are similar, both being accentuated towards walls, corners and cues. Note the local increase at cue boundaries observed for both visual and population change (black lines in **C** indicate location of wall mounted cues). (**G**) Population activity change correlates more strongly with visual change (r = 0.60) than speed (r = -0.30), turning rate (r = 0.27) and distance to the nearest walls orthogonal (r = -0.17) or parallel (r = 0.05) to the rats’ motion. (**H**) Filtering by heading direction reveals how the change in both the visual scene and population activity depends on proximity to walls and their orientation relative to the direction of travel.

Finally, to investigate if visual information contributed to this effect, we developed a VR replica of environment D (Fig. S12). A 300° field of view was used to reconstruct each animals’ visual scene at 1 cm increments along its trajectory (Fig. 4C), and the visual change between consecutive frames was calculated during motion (speed >10 cm/s, see Methods). We found a tight coupling between the change in visual scene and change in place cell population activity (Fig. 4D&G; time series correlation: Pearson’s r=0.60, p < 0.001, N = 510871) - a stronger relationship than was found for other behavioural variables (Fig. 4G) and which persisted when they were accounted for (partial correlation: r=0.51, p < 0.001, N = 510871). As expected, when mapped out in space, both the activity vector change and visual change showed a general increase with proximity to the walls, corners and the two wall mounted cues (Fig. 4E&F; no smoothing applied), as well as a strong correlation between both maps (correlation between maps: Pearson’s r=0.72, p < 0.001, N = 5524). Importantly, filtering the behaviour by heading direction (Fig. 4H) emphasises how this variation depends not only on position but also on movement direction, with walls and cues orthogonal to the direction of motion being key factors in driving the visual change - a pattern that closely mirrored the direction-filtered activity vector change (all Pearson’s r > 0.66, all p < 0.001, N > 5306).

## Discussion

Using high-yield recordings from rodents foraging in large environments, we have shown that while the distribution and extent of individual place fields are governed by proximity to environmental boundaries, the statistics of the population as a whole are effectively constant. Specifically, the widths of place fields displayed a Weber’s law-like (Fechner, 1860) increase with distance to the orthogonal wall, which was exactly opposed by a commensurate decrease in field count. Thus, in general small place fields were densely distributed near the boundaries of the environments, while broader fields were found more sparsely towards the middle of the open spaces. Therefore, the rate of change in the activity of the population depends on both boundary proximity and movement direction, in a manner that correlates strongly with the rate of change in the animals’ reconstructed visual scenes.

Our results encapsulate prior findings that individual cells have more and larger fields in bigger spaces (Fenton et al., 2008; Rich et al., 2014), describing them in terms of a simple interaction between homeostatic balance at the population level and rate of perceptual change, which is naturally elevated near boundaries and cues. Equally, this same relationship accounts for the clustering of place fields around visual cues in a 1D VR (Bourboulou et al., 2019) and the effect of manipulated visual gain on hippocampal place cells in a 1D and 2D VR (Campbell et al., 2018; Chen et al., 2019). This work also resonates with early geometric cue-based models of place cell activity which sought to describe place fields using Weber-like responses to environmental boundaries (Barry et al., 2006; Hartley et al., 2000; O’Keefe and Burgess, 1996). Notably, these models focused on the description of individual fields but were agnostic of population-level interactions which featured in models that emphasised CA3 recurrence and mutual inhibition (Káli and Dayan, 2000; Samsonovich and McNaughton, 1997). Thus, our current results indicate that a synthesis of both approaches is important for understanding how the hippocampus represents large-scale spaces - yoking the evolution of statistically stable population-level activity to movement through visual states (Hedrick and Zhang, 2016; Uria et al., 2020).

The precise mechanism that maintains the observed population-level statistics is unclear but might plausibly result from metabolic homeostasis imposed by limitations on blood flow, or the energetic limitations of the neurons themselves (Laughlin et al., 1998), and could be modelled as competitive learning between hippocampal inputs for these limited resources (Barry and Burgess, 2007). Equally, it may well be a natural limitation of the highly recurrent CA3 network and the need to balance excitation with inhibition to avoid a runaway increase in activity (Hasselmo et al., 1995). Either way, the highly stable population-level statistics present an interesting conundrum. In a stationary animal, there is no statistical difference in the composition of hippocampal activity between large or small environments or between different positions in those environments. Thus, without decoding the precise location it does not appear to be possible to make any inference about the general characteristics of the animal’s current location. In contrast, during movement the rate at which the population evolves relative to the animal’s velocity does provide some information about the current setting but even that is dependent on the direction of travel. In short it appears there is no simple capacity for a high-level meta-code capable of describing common elements of different environments. Instead, the place cell map may develop randomly within homeostatic constraints, its resolution being determined by the granularity of the animal’s sensory environment.

Importantly, the relationship between visual change and place cell fidelity can be understood in terms of Information theory. The rate of change of perceptual states is tightly linked to the amount of Fisher information they transmit about an unknown parameter (Wei and Stocker, 2017), in this case position. Regions where the visual scene changes rapidly with respect to position - near the walls for example - convey more information about self-location than regions where visual stimuli change more slowly. In turn, for simple neural codes in one and two dimensions, the tuning width of a given neuron is inversely proportional to the Fisher information it carries (Brown and Bäcker, 2006; Zhang and Sejnowski, 1999). If we make the reasonable assumption that the place code maximises information transmitted up to the limit imposed by vision, then this directly predicts that place field width will be inversely proportional to the rate of change in the visual scene. Alternatively stated, rate of change in place cell population activity is expected to be proportional to visual change - the result we observe. Notably this relationship is not necessarily specific to vision or place cells. Thus, it is likely that place cell fidelity will also be modulated by information conveyed in different sensory modalities, such as tactile changes and olfactory gradients. More generally, it seems plausible that the scale and fidelity of other neural representations of self-location must be subject to the same Information theoretic limits. Indeed, the increase in entorhinal grid cell scale noted towards the centre of large environments (Hägglund et al., 2019), and potentially other spatial distortions (Krupic et al., 2015; Stensola et al., 2015) can be seen through the same lens.

## Methods

### Animals and tetrode implantation

Five male Lister Hooded rats were used for this study. All procedures were approved by the UK Home Office, subject to the restrictions and provisions contained in the Animals Scientific Procedures Act of 1986. All rats (333-386 g/13-17 weeks old at implantation) were implanted with two microdrives targeted to the right and left CA1 (ML: 2.5 mm, AP: 3.8 mm posterior to bregma, DV: 1.6 mm from dura) following a standard surgery and recovery procedure (Barry et al., 2007). After surgery, rats were housed individually in Perspex cages (70 cm long x 45 cm wide x 30 cm high) on a 12 hr light/dark cycle. Screening and experiments took place during the dark phase of the cycle. After one week of recovery, rats were maintained at 90-95% of free-feeding weight with ad libitum access to water.

The hair around the incision site was removed, and the skin was sterilised with Betadine. The animal was placed on a heating pad for the duration of the surgery to maintain body temperature. Viscotears Liquid Gel was used to protect the animal’s eyes. General anaesthesia during the operation was maintained with an isoflurane-oxygen mix of 1.5-3% at 3 l/min. Carprieve (1:10) and Baytril were injected subcutaneously (0.1 ml/100 g) before the surgery for analgesia and to minimize chances of infection, respectively. Baytril was also included in post- operative treatment in their water for one week. An analgesic, Metacam Oral Suspension suspended in jelly, was administered for three days post-surgery. The tetrodes were implanted through ∼1 mm trephine craniotomies over target sites, and they were fixed to the exposed skull with dental cement (Super-Bond C&B) and six bone screws. A gold pin used as ground and reference was soldered to one of the orbital bone screws before its implantation. The craniotomies and elements of the microdrives were protected from dental cement using Vaseline.

### Electrophysiological and behavioural recordings

Each single-screw microdrive (Axona Ltd.) was assembled with two 32 channel Omnetics connectors (A79026-001), 16 tetrodes of twisted wires (either 17 µm H HL coated platinum-iridium, 90% and 10% respectively, or 12.7 µm HM-L coated Stablohm 650; California Fine Wire), and platinum-plated to reduce impedance to below 150 kΩ at 1 kHz (NanoZ).

Electrophysiological recordings were acquired using Open Ephys recording system (Siegle et al., 2017) and a 64-channel amplifier board per drive (Intan RHD2164). The recorded signal was referenced to an orbital bone screw - also the ground for the amplifier boards. The Open Ephys Acquisition Board was grounded to an aluminium foil sheet positioned underneath the vinyl flooring throughout the entire extent of the experimental room. Electrophysiological signals were recorded from 128 channels at 30 kHz. Spikes were detected as negative threshold crossings of more than 50 µV in the 30 kHz signal after bandpass filtering between 600 and 6000 Hz. For each spike, waveforms were stored at 30 kHz for the 1.2 ms window surrounding the threshold crossing. The waveforms are displayed and discussed in their inverted form, where the largest deflection from baseline is a positive peak.

Positional tracking was performed with an open-source multi-camera tracking system SpatialAutoDACQ(Tanni, Sander and Barry, Caswell, 2021). The output position data from SpatialAutoDACQ was the spatial coordinate of an infra-red LED positioned above the animal’s ears, sampled at 30 Hz.

### Histology

Anatomical locations of recordings were verified using histology. Rats were anaesthetised with isoflurane and given intraperitoneal injection of Euthatal (sodium pentobarbital) overdose (0.5 ml / 100 g) after which they were transcardially perfused with saline, followed by a 10% Formalin solution. Brains were removed and stored in 10% Formalin and 30% sucrose solution for 3-4 days before sectioning. Subsequently, 50 µm frozen coronal sections were cut using a cryostat, mounted on gelatine coated or positively charged glass slides, stained with cresyl violet and cleared with clearing agent (Histo-Clear II), before covering with DPX and coverslips. Sections were then inspected using an Olympus microscope, and tetrode tracks reaching into CA1 pyramidal cell layer were verified.

### Experimental paradigm

Screenings for a suitable place cell yield were performed from one week after surgery in a 1.4 x 1.4 m environment, different from those used in any of the experiments. Tetrodes were gradually advanced in 62.5 µm steps until ripple oscillations could be observed, and pyramidal cells with stable firing fields could be identified.

Screenings, training, and experiments all took place in the same experimental room, using environments constructed of the same materials. Environments had black vinyl flooring; were constructed of 60 cm high modular boundaries (MDF) coloured matt black, surrounded by black curtains on the sides and above. Each environment was illuminated by an elevated (2 m) diffuse daylight lamp from each corner of the environment that was adjacent to a corner of the experimental room (multiple lights in larger environments), with each lamp producing between 30-50 Lux/m. All experiments involved scattered 20 mg chocolate-flavoured pellets (Dustless Precision Pellets® Rodent, Purified, Bio-Serv, USA) dropped into the environment by an automated system in SpatialAutoDACQ to encourage foraging. The automated system scattered the pellets randomly with greater preference for areas least visited by the animal.

Place cells were recorded as each animal foraged in four environments of different size (Fig. 1A, Fig. S1). The environment sizes were 87.5 x 125 cm (environment A), 175 x 125 cm (environment B), 175 x 250 cm (environment C) and 350 x 250 cm (environment D). All environments were rectangles with close to identical shape (axes ratio 1.40). Environment A was the smallest, and the other sequentially larger environments – B, C and D – each doubled in size by doubling the length of the shortest axis (Fig. 1A). There were two sets of cues in the environments. The most prominent cue elevated above the wall of the enclosure was different in all environments, varying in size and the type of pattern, but always black and white. Two different secondary smaller cues were used, an A4 sheet (11 × 16 cm) with a dot pattern and a set of three adjacent A4 pages, placed on the wall at a height the animal could not reach. The number of secondary cues and the size of the primary cues varied slightly between environments to scale with their size (Fig. 1A).

During a single session consisting of 5 trials, the place cells of an animal were recorded in all four environments and twice in environment A (Fig. 1B). The second recording in environment A is referred to as a recording in environment A’. The duration of the recording in the smallest environment (A) was 15 minutes. The recording duration doubled along with the size of each environment, reaching 120 minutes in the largest environment (D). Each animal was recorded on three or four sessions, the data analysed here comes from the first session in which an animal achieved good spatial sampling in all environments (sessions 3, 2, 2, 3, and 4, for the 5 animals).

After a recording in each environment, the flooring was wiped with an unscented soap solution to clear any potential olfactory cues. The animal was kept in a familiar rest box (with water provided) between each recording for 15 to 30 minutes, while the preceding and following environments were disassembled and reassembled, respectively.

### Cell identification

Spikes were assigned unit identities with automated clustering software (KlustaKwik) (Kadir et al., 2014) based on spike waveforms. The results from the automated clustering were curated using an offline data analysis suite (Tint, Axona Ltd., St. Albans, UK) to further separate under-clustered units and merge over-clustered units.

All recordings from a given animal that were performed on the same session were clustered simultaneously, concatenating the spike waveform data. Therefore, the same set of units were identified across all such recordings. This approach made it possible to analyse the properties of the same place cell population in multiple conditions.

L-ratio (Schmitzer-Torbert and Redish, 2004) and Isolation Distance (Harris et al., 2001) were calculated through Mahalanobis distance to verify that sorting quality has not affected the results. The features used for this analysis were the same as those used for automated and manual clustering: amplitude, time-to-peak, time-to-trough, peak-to-trough, half-width, trough- ratio, and the first three PCA components of waveforms. These measures were computed on waveforms combining all clusters, including noise, and pooling across all recordings – the same way as was done for spike sorting.

Place cells were identified computationally after the clustering procedure. The following criteria were used to identify place cells:

- Waveform peak-to-trough duration of over 0.45 ms.
- Waveform peak half-width of over 0.1 ms.
- The ratio between amplitude and trough voltage values (trough-ratio) of over 0.175.
- Spatial correlation of odd and even minute ratemaps of over 0.5 in at least one recording.
- Spatial correlation of first and last half ratemaps of over 0.25 in at least one recording.
- At least one field in one of the recordings (field detection method described below).
- Mean firing rate across all recordings lower than 4 Hz.

Place cells were further filtered for duplicates recorded on separate tetrodes. Duplicate units were considered to be unit pairs that passed the following criteria mostly based on cross-correlograms with 2 ms bins and a maximum lag of 25 ms:

- At least 200 spikes at 0-lag.
- Lower than 0.5 ms sigma of a gaussian fitted to the cross-correlogram.
- Mean spatial correlation of ratemaps higher than 0.5 across recordings where both units have at least 200 spikes.

If duplicate units were detected, the one with more total spikes was set as noise, to maximise signal to noise ratio. This approach was based on the observation that the unit in the duplicated pair that had more spikes was usually less well isolated from noise or other units.

Interneurons were identified based on the following criteria:

- Minimum mean firing rate of 4 Hz across all recordings.
- Maximum waveform half-width of 150 μs.
- Maximum trough-ratio of 0.4.
- Maximum spatial correlation of 0.75 in any environment.

### Computing ratemaps

To calculate a ratemap for each unit, the position data was binned into 4 cm square bins, and the number of position samples in each bin was divided by the sampling rate, producing the dwell time for each spatial bin. Spike timestamps were paired with simultaneous position samples and assigned to corresponding spatial bins, thereby producing spike counts for each spatial bin. Only the position samples and spike timestamps from periods where the animal was moving at more than 10 cm/s were used to produce these dwell time and spike count maps. Both dwell times and spike counts were smoothed with a Gaussian kernel (standard deviation of 2 spatial bins) while setting unsampled bin values and those outside the environment to 0. The resulting smoothed spike counts were divided by smoothed dwell times, producing spatial ratemaps.

### Place field detection

Place fields were detected in spatial ratemaps to analyse place cell properties at the level of individual place fields. Here, a place field is defined as a contiguous area in a ratemap, where the firing rate decays continuously from a single prominent peak, and the observed firing rates are well correlated across multiple visits to the same location. Often individual place fields are so close to each other that the firing rate threshold traditionally used for place field detection (1 Hz) would not be able to detect them as separate place fields. This effect is exacerbated by the spatial smoothing step in computing spatial ratemaps. However, based on the definition above, these areas should be considered as separate place fields.

An iterative thresholding method was used to find spatial bins that constituted a single place field in a ratemap (Fig. S2). As a first step, the ratemap was thresholded at 1 Hz, and contiguous groups of bins (ignoring diagonal connections) were identified as candidate fields. The regions including at least 10 spatial bins and a peak value of at least 2 Hz were considered as valid candidate fields.

The ratemap of each valid candidate field was then thresholded again with a 0.05 Hz higher threshold (1.05 Hz), and the same method of finding contiguous regions and their validation was applied. This was done iteratively, resulting in continuously smaller regions, each with a higher threshold and associated with their parent field candidates with a lower threshold, some having more than one child field candidate.

The resulting lists were then parsed in reverse order, starting with the smallest candidate fields with highest thresholds. Candidate fields that were too large (greater than half the bins of the ratemap) or not sufficiently stable over repeated visits to the region (spatial correlation of odd and even minute ratemaps below 0.25) were ignored. As the increasingly lower threshold candidate fields overlapping with each other were assessed, the lowest threshold valid candidate field in a sequence of overlapping candidate fields was detected as a place field. The overlapping candidate fields with a higher threshold were ignored. If more than one child candidate field of a lower threshold candidate field was valid, the large single candidate field was ignored, and the smaller valid candidate fields were detected as separate place fields. In this manner, multiple place fields were detected in individual ratemaps of single units, as illustrated in Fig. S2.

### Statistical methods

In most cases Kruskal-Wallis test was used to test for differences between groups due to small sample sizes. Positive Kruskal-Wallis tests of more than two samples were followed by two-sided

Mann-Whitney U test for individual pairwise comparisons, and Benjamini/Hochberg (non-negative) correction (Benjamini and Hochberg, 1995) for multiple comparisons. Benjamini/Hochberg (non-negative) correction was implemented in Statsmodels Python package (Seabold and Perktold, 2010) and Kruskal-Wallis, Mann-Whitney U, Linear regression, Pearson coefficients, Poisson and Gamma distributions were computed using Python statistics package Scipy (Virtanen et al., 2020).

### Position decoding

Position decoding was used to estimate the location encoded in the activity of a place cell population at specific timepoints. The probability of the animal being at each location in the environment is computed based on the similarity between the ongoing firing rates of place cells in a time-window (e.g. 1 second) and their spatial ratemaps. The decoded location is then identified as the one with the highest likelihood. The spatial ratemaps for this purpose were computed using periods where the animal was moving faster than 10 cm/s, excluding the time- point that was being decoded – cross-validation with a 3-minute window. The method of matching the ongoing population activity to spatial ratemaps has been used previously (Mathis et al., 2012; Towse et al., 2014) and is based on the original formulation by Zhang et al. (1998).

Specifically, the population activity of *N* units *K* = (*k*_1_, . . . . , *k*_*N*_) was computed, where *k*_*i*_ is the spike rate of the i-th unit in a temporal bin (e.g. 1 second). Expected population activity *a* at location *x*, belonging to the set of all spatial ratemap bin centres *X*, was based on the values of all units in the spatial ratemap corresponding to that location bin *a*(*x*) = (*f*_*i*_, . . . , *f*_*M*_), such that *a*_*i*_(*x*) = *f*_*i*_, refers to the value in the spatial ratemap of unit *i* at location *x*. These representations of neural activity were used to compute the conditional probability of observing *K*, at location *x* as:

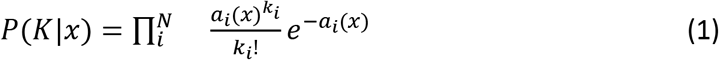

This method allows assessing the probability of any spatial bin being decoded independently of the number of bins considered and their spatial arrangement. It is agnostic to the animal’s real location and past decoded locations, as it considers all locations to have equal prior probability – it has a flat prior. The location encoded in the population activity *x*^(*K*) was then computed as the centre of the spatial bin with the highest conditional probability:

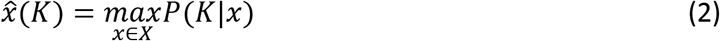

To decode the environment from the place cell population activity (Fig. S3), the posterior probability distribution (equation 1) was calculated over all visited bins in all environments. The environment pertaining to the most probable spatial bin (equation 2) was then identified as the decoded environment.

### Spatial correlation

Spatial correlation was used to quantify the similarity between spatial ratemaps of pairs of cells or spatial ratemaps of the same cell that were constructed using data from different parts of the same recording. Spatial correlation was the Pearson correlation coefficient for pairs of values from spatial bins with matching locations in two ratemaps. For a spatial bin to be included, it must have had a firing rate above 0.01 Hz in at least one of the ratemaps to avoid high correlations between 0 Hz bins. Unvisited bins were ignored. At least 6 such valid spatial bins were required for spatial correlation to be computed, which was always the case for place cells.

### Field formation models

Two models were used to estimate place field count per cell as a function of environment size: equal-Poisson and gamma-Poisson models. The equal-Poisson model has one parameter, the average field formation propensity *τ*, which is constant for all cells, and the model predicts the field counts per cell *X* as a function of *τ* and environment area in m^2^ *A*

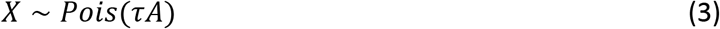

The gamma-Poisson model estimates the field formation propensities *T* in the place cell population based on a shape *α* and scale *θA*parameters

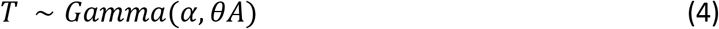

These field formation propensities *T* are then used to estimate the place field counts *X*

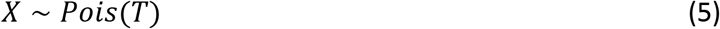

Gamma-Poisson can then be defined using a negative binomial

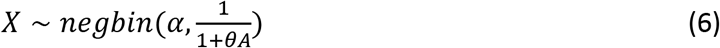

Using the change of variables 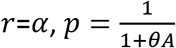

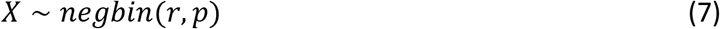

giving the gamma-Poisson probability mass function

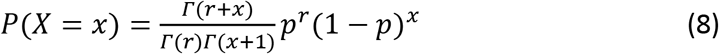

The parameters for both models were optimised using maximum likelihood estimation using the field counts of all place cells (N = 627) in the combined environment - counting fields per cell across all four environments. The parameter optimisation was performed with L-BFGS-B solver implemented in the Scipy Python package (Virtanen et al., 2020). The two models were compared using the Bayesian information criterion.

The gamma-Poisson model was fit as described above, and then evaluated on prediction of proportion of cell recruitment with the physical environment size values and also with environment sizes adjusted based on field density. The latter always performed better, therefore, all the reported results were computed using environment sizes adjusted based on field density. The proportion of fields in each environment was modelled with linear regression *y* = *bA* + *c*, where *A* is the area of an environment (Fig. 2D). Therefore, using the same parameters *b* and *c* the field density *ρ* in an environment of size *A* can be computed as

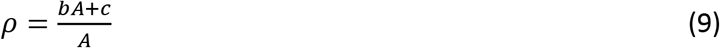

The field density adjusted environment size *A*′ for computing the gamma-Poisson probability mass function in an environment with size *A* was computed as

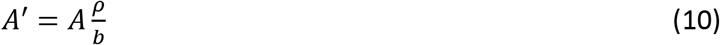

Proportion of place cells recruited to form at least one field in the four environments (A, B, C and D) based on the gamma-Poisson model was computed by modelling a population of 100,000 cells. Each cell had a field formation propensity *τ* (drawn from the gamma distribution defined by *α* and *θ* fit already previously to field formation propensities) that was used to compute the number of fields using a Poisson process with rate *τA* or *τA*′. Modelled cells with no field in any of the environments were ignored to match the conditions applied to the experimental data.

### Field size measures

The place field areas were computed by summing the area of all spatial bins (16 cm^2^) covered by each detected field. The place field widths in the two axes were computed as the length of the field’s projection onto a given axis (Fig. S7). These values were used to calculate the mean field area and width in each animal at every spatial bin by averaging the values of all fields overlapping a particular spatial bin. To estimate the average field area and width at different distances to the wall, the values for spatial bins in a particular range of distance from the wall (e.g. 0-25 cm) were averaged separately for each animal. Where further spatial selection is indicated in the cartoons above figures (e.g. only including data from the middle third of the environment), the averaged spatial bins were selected in such manner to minimize the effects from orthogonal walls.

### Population activity summary statistics

The proportion of co-active place cells and the mean firing rate of place cells at different distances to the wall were computed by averaging the spatial ratemap values in all spatial bins that were in that range of distances from the wall. The proportion of co-active cells (firing rate ≥1 Hz) and the mean firing rate was computed using the spatial ratemap values at a given location, including all place cells detected in a given animal.

The population firing rate distributions were computed by aggregating the Gaussian smoothed (1 second sigma) firing rates aligned to position samples (30 Hz) where the animal’s location was within a particular range of distance from the wall. At each timepoint, only the activity of neurons with firing rate ≥1 Hz were included to facilitate measuring firing rate distributions. Only samples when the animal was moving faster than 10 cm/s were included. The samples assigned to each range of distances from the wall were further split temporally into the first and second half. The Jensen-Shannon divergence was then computed between each pair of halves, and also between all halves at different distances to the wall, to compare the divergence due to inherent variation and location.

### Population activity change

Position data smoothed with Savitzky–Golay filter (166 ms window and polynomial order 5) and place cell firing rates computed at 33 ms bins and smoothed with a Gaussian (166 ms sigma) were used to construct population activity vectors for computing the population activity change. The position data was reduced to one dimension by computing the Euclidean distance cumulatively over consecutive samples. It was then used to linearly interpolate cell firing rates to position samples 1 cm apart. The population activity change was then computed as the Euclidean distance between consecutive samples of place cell firing rates - population activity vectors measured at 1 cm intervals. All place cells detected in an animal were included and periods where the animal was moving slower than 10 cm/s were excluded.

### Visual change

The virtual environment was created in Unity3D with the same proportions as the physical environment. Animal trajectories were speed filtered (>10 cm/s) and interpolated so that consecutive samples were 1 cm apart (equivalent to method in population activity change). The visual scene from each sample point was then captured by three greyscale cameras, raised the equivalent of 5 cm from the floor and angled 35° above the horizontal axis. These cameras were oriented 100° apart in the horizontal plane, and each rendered a 64×64 pixel image with a field of view of 100° to give a total field of view of 300°. The absolute difference between consecutively sampled greyscale images was used to yield the pixel-by-pixel change at each sampling point. This pixel-by-pixel change was then z-scored per pixel and averaged across pixels to generate a single value for visual change at each sample point.

Visual change and population activity change maps represent the average value of samples across the environment using a 4×4 cm bin size and no smoothing applied. For illustration purposes, the time series presented in Fig. 4D was smoothed with a 1D boxcar filter of width 3, but no smoothing was applied when calculating the reported correlations between time series.

## Acknowledgements

We would like to thank Tim Behrens, Dan Bush, Julie Lefort and Romain Bourboulou for useful comments on the manuscript.

The authors S.T., W.d.C. and C.B. thank the Wellcome Trust for supporting this work through the Senior Research Fellowship awarded to C.B. (212281/Z/18/Z) and the Medical Research Council UK for the PhD studentship awarded to S.T.

This research was funded in whole, or in part, by the Wellcome Trust [212281/Z/18/Z]. For the purpose of open access, the author has applied a CC BY public copyright licence to any Author Accepted Manuscript version arising from this submission.

## Author Contributions

S.T. and C.B. conceptualized the experiment. S.T. collected and spike-sorted the data. S.T. and C.B. analysed the electrophysiology and behavioural data. W.d.C. created the virtual replica, performed the simulations and related analysis. S.T., W.d.C., and C.B. wrote the manuscript.

## Data availability

The data that support the findings of this study are available from the corresponding author upon reasonable request.

## Code availability

Code for data collection is available at https://barry-lab.github.io/SpatialAutoDACQ/

Code for analysis is available at https://github.com/Barry-lab/Publication_TanniDeCothiBarry2021

## Supplementary Information

**Fig. S1.**
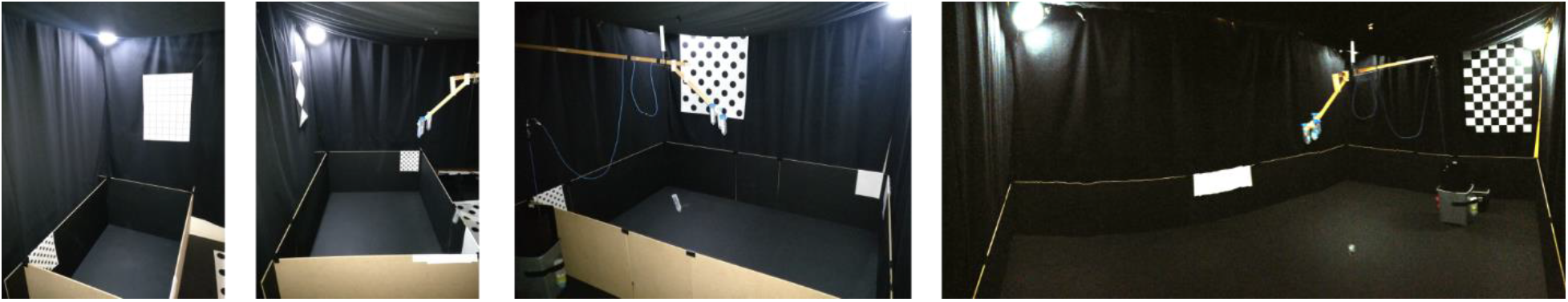
Photographs of experimental environments. Environments from left: A, B, C and D.

**Fig. S2.**
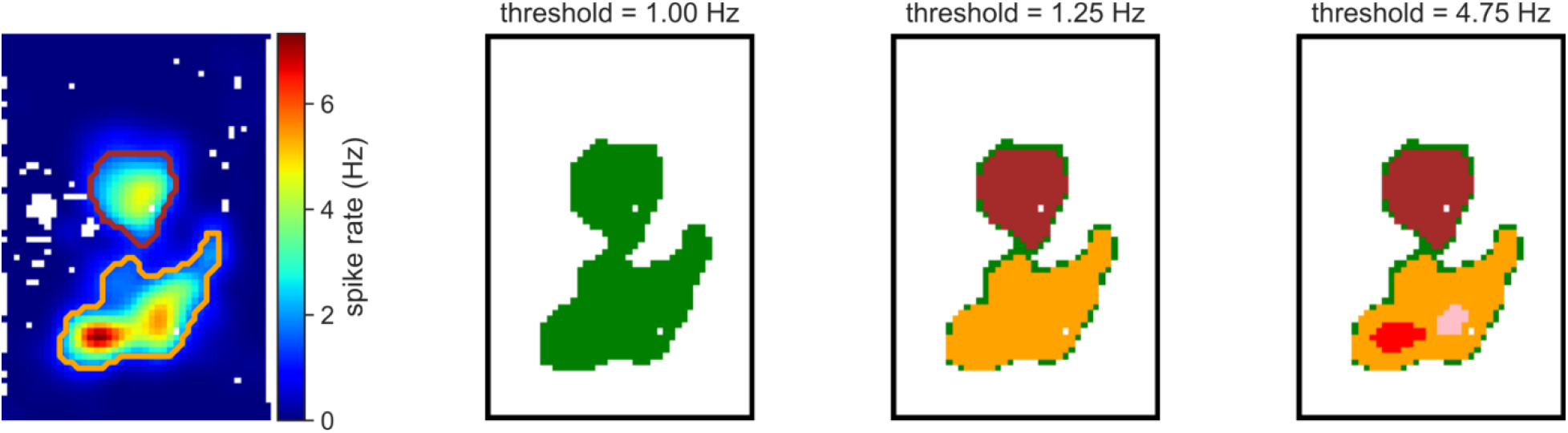
Field detection method based on iterative thresholding identifies multiple place fields in a single ratemap. The spatial ratemap of a place cell is shown on the left. With the threshold at 1 Hz, only one place field (green) is detected. While increasing the threshold at 0.05 Hz increments, two place fields (brown and orange) are identified with 1.25 Hz threshold. These both pass the place field criteria. By increasing the threshold further, two smaller place fields (red and pink) are detected with a 4.75 Hz threshold, both overlapping with the larger orange place field. At least one of the smaller place fields (red and pink) did not pass the place field criteria. Therefore, both of them (red and pink) were ignored because a larger place field, detected with a lower threshold and overlapping with them, did pass the place field criteria.

**Fig. S3.**
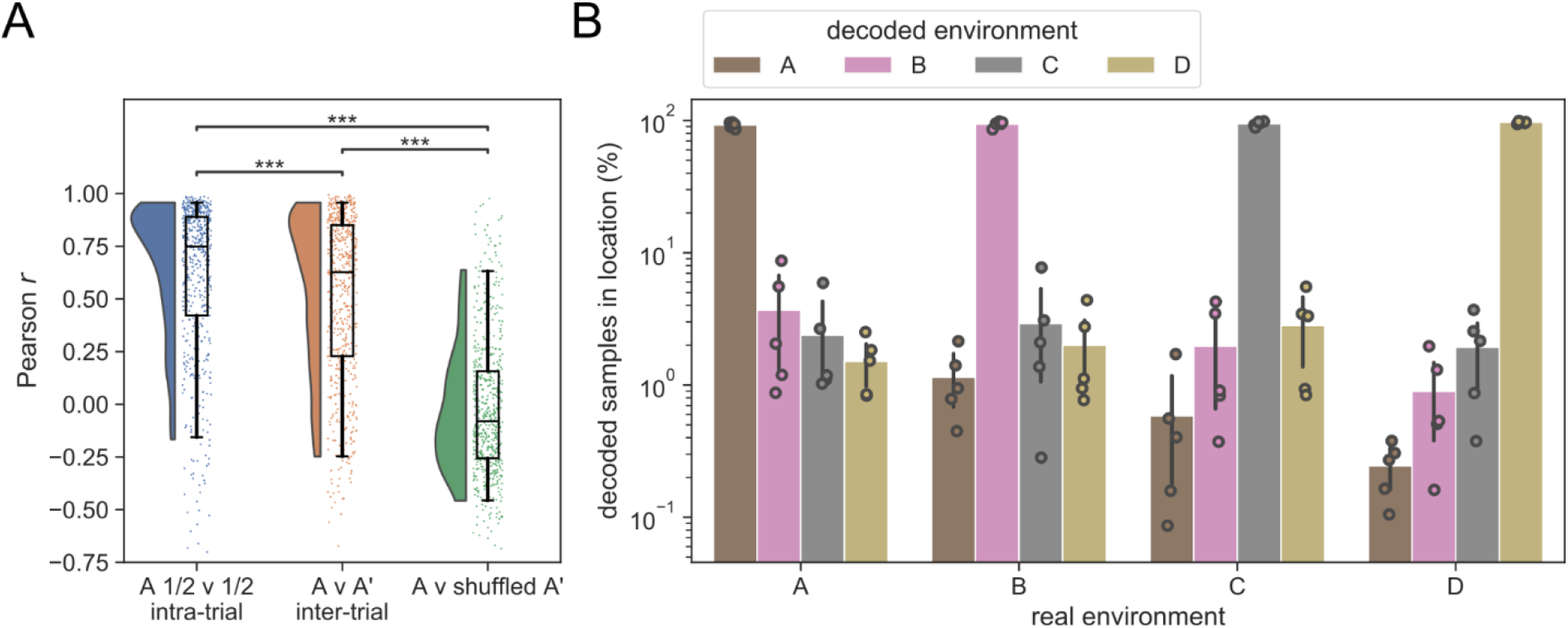
Place cells formed distinct, stable representations for each environment. (**A**) Spatial correlations (bin-wise Pearson correlation) between rate maps from the first and second half of environment A (A ½ v ½) and between repeated trials (A v A’) were high (mean spatial correlation of 0.61 and 0.51, respectively), significantly exceeding the values obtained by randomly repairing cells (A v shuffled A’) (-0.04). Kruskal-Wallis test: H = 644, p = 1.64 x 10^-140^; Mann-Whitney for A ½ v ½ and A v A’, U = 1.8 x 10^5^, p = 3.2 x 10^-6^, for A ½ v ½ and A v shuffled A’, U = 2.8 x 10^5^, p = 10^-92^, for A v A’ and A v shuffled A’, U = 2.9 x 10^5^, p = 10^-114^. The box shows quartiles of the dataset, and whiskers indicate the 5th and 95th percentile of the data distribution. The kernel-density estimate is bounded between the 5th and 95th percentile. (**B**) Population activity vectors were reliably decoded to the environment from which they were drawn - Bayesian-framework with 1s window used for decoding. Error bars show 95% confidence intervals of the mean based on bootstrapping.

**Fig. S4.**
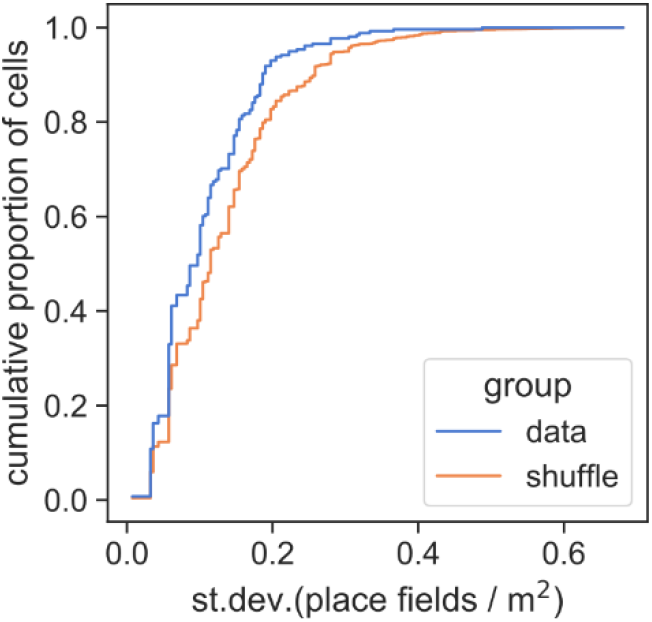
Place cell field formation propensity is conserved across environments. The standard deviation of place cells’ field formation propensity (place fields / m^2^, accounting for field density in Fig. 2D inset) across environments was lower than for a shuffled distribution (Mann-Whitney: U = 2.7 x 10^7^; p = 6 x 10^-8^; N = 258). Only cells with at least 1 field in each environment were used in this analysis. Shuffle was obtained by permuting cell identities within each animal and environment 1000 times.

**Fig. S5.**
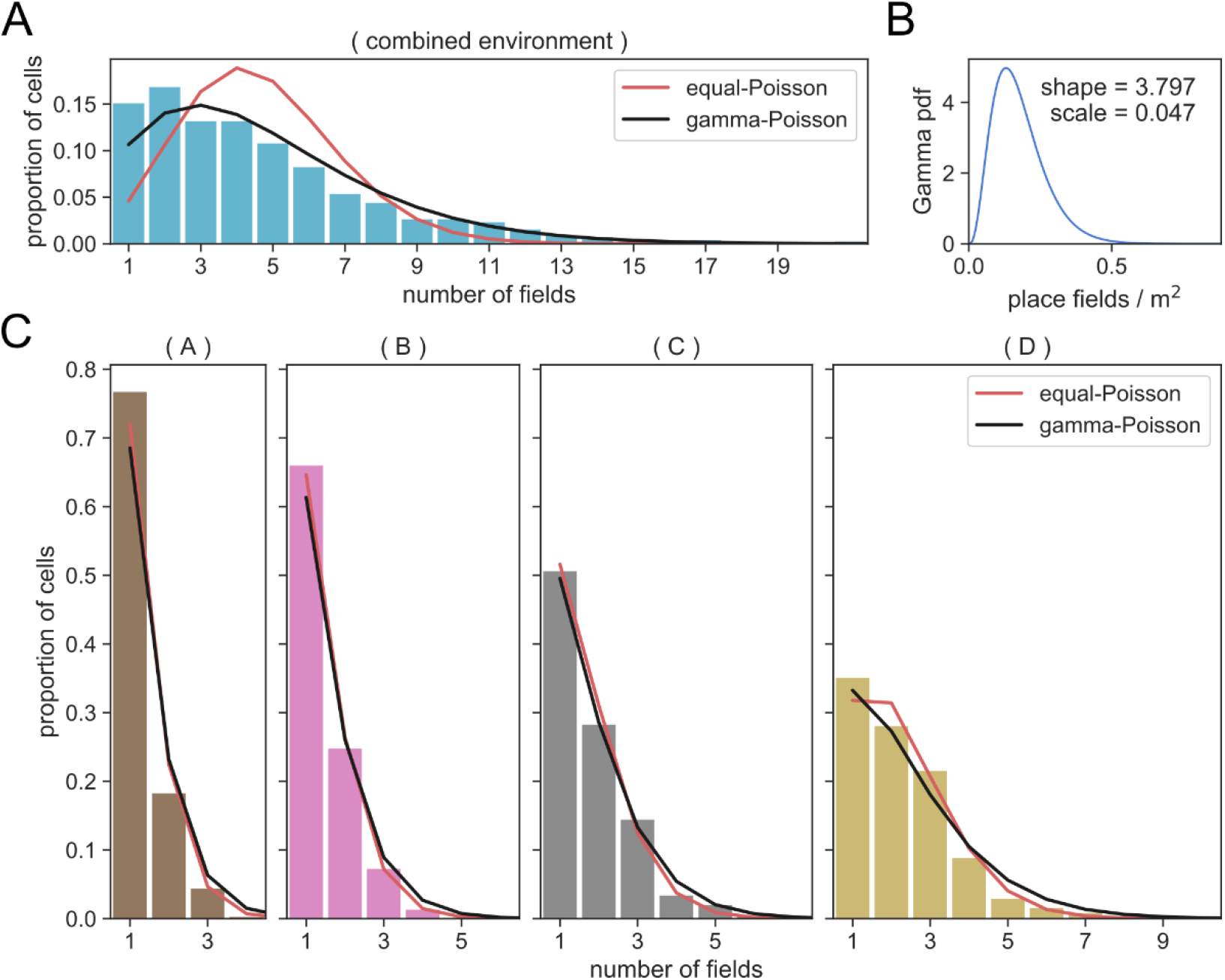
Place field formation propensities and the gamma-Poisson model. (**A**) Distribution of field counts per place cell after grouping all environments together and predictions of the two models fit to this data. (**B**) Probability density function (pdf) of gamma with fitted parameters, defining the field propensity distribution as a function of environment size. (**C**) Distribution of field counts per place cell for cells in each environment that have at least one place field, and predictions of the two models fit to data in **A**.

**Fig. S6.**
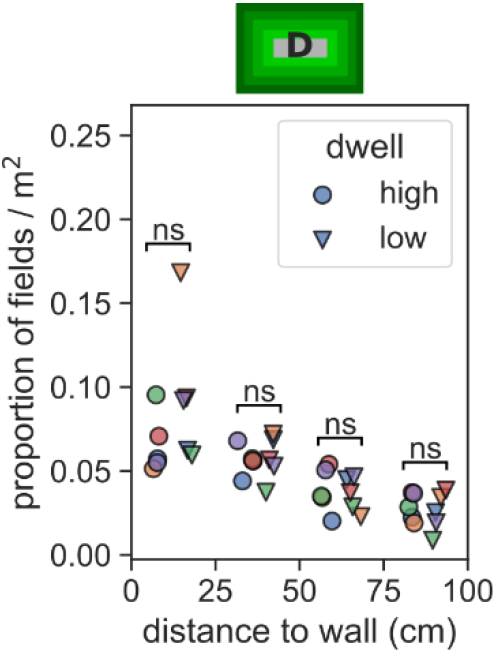
Change in field density with distance to wall is not explained by the difference in dwell time. Field density (proportion of fields / m^2^) and dwell time were computed for 25 x 25 cm non-overlapping regions of the largest environment chosen to be at a range of distances to the nearest wall. For each animal, we grouped regions according to distance to the wall and for each of these groups found the mean field density in the region with the highest and lowest dwell time. Field density did not vary with dwell time (Mann-Whitney for all comparisons: U ≥ 6, p ≥ 0.11) but was different between regions at different distances to the wall (Kruskal-Wallis for both high and low dwell: H ≥ 13.1, p ≤ 0.004).

**Fig. S7.**
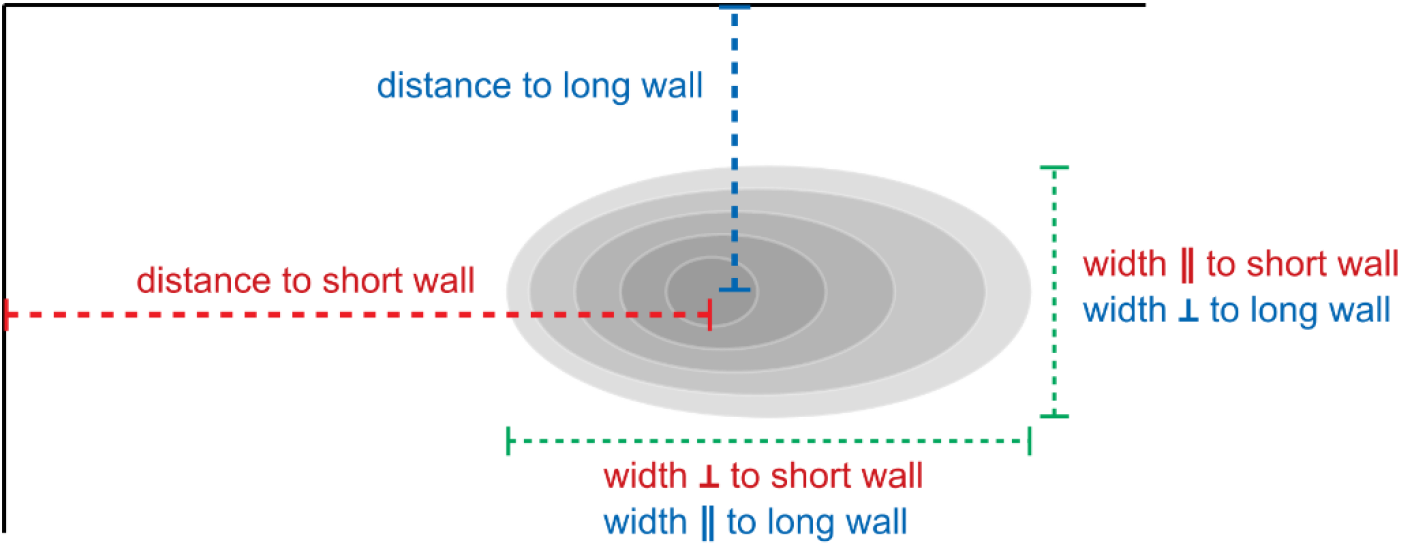
Place field width measurements. The width of each place field was measured along two orthogonal axes. The distance of the place field from nearest walls in the two axes was measured from the location of peak firing rate.

**Fig. S8.**
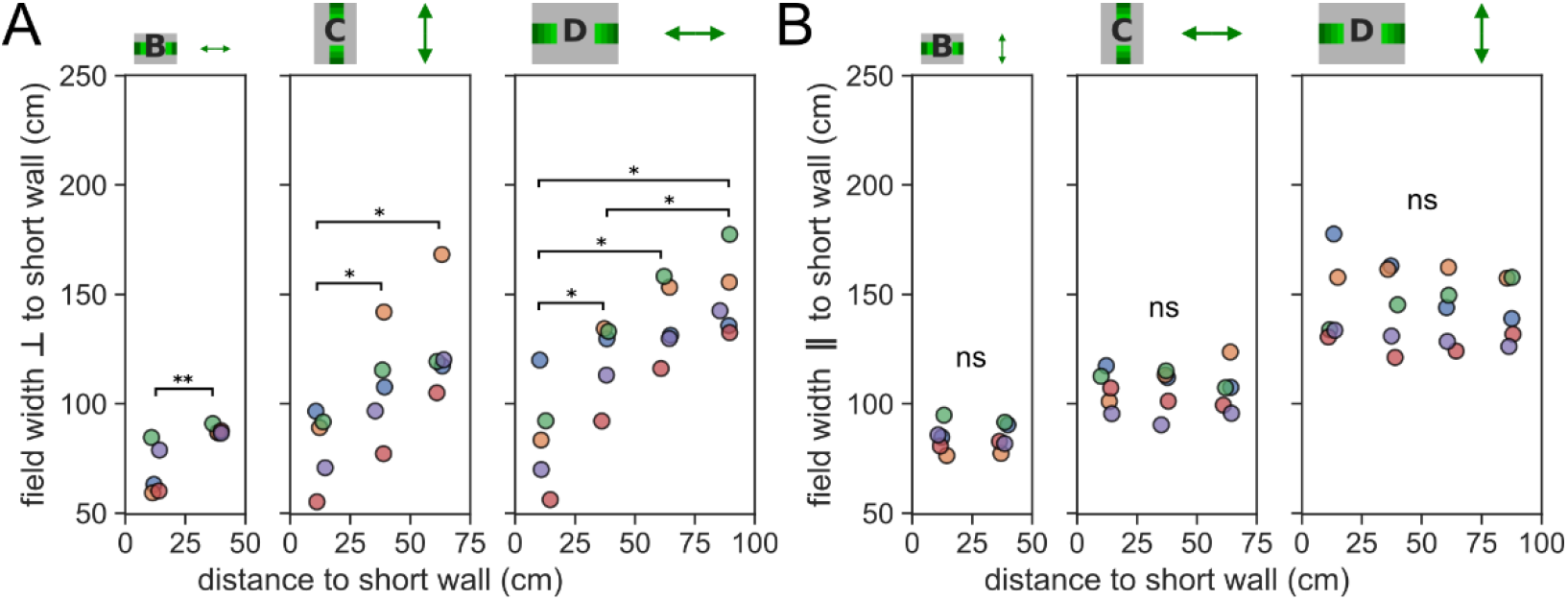
Field width orthogonal to a nearby wall increases with distance to that wall in all environments. (**A**) The average place field size per animal measured orthogonal to the short wall increases with distance from the wall and is also greater near the wall in the larger environments. Pair-wise post hoc tests adjusted for multiple comparisons using Benjamini/Hochberg (non-negative) correction. (**B**) The average place field size per animal measured parallel to the short wall is constant at all distances to the short wall in all environments.

**Fig. S9.**
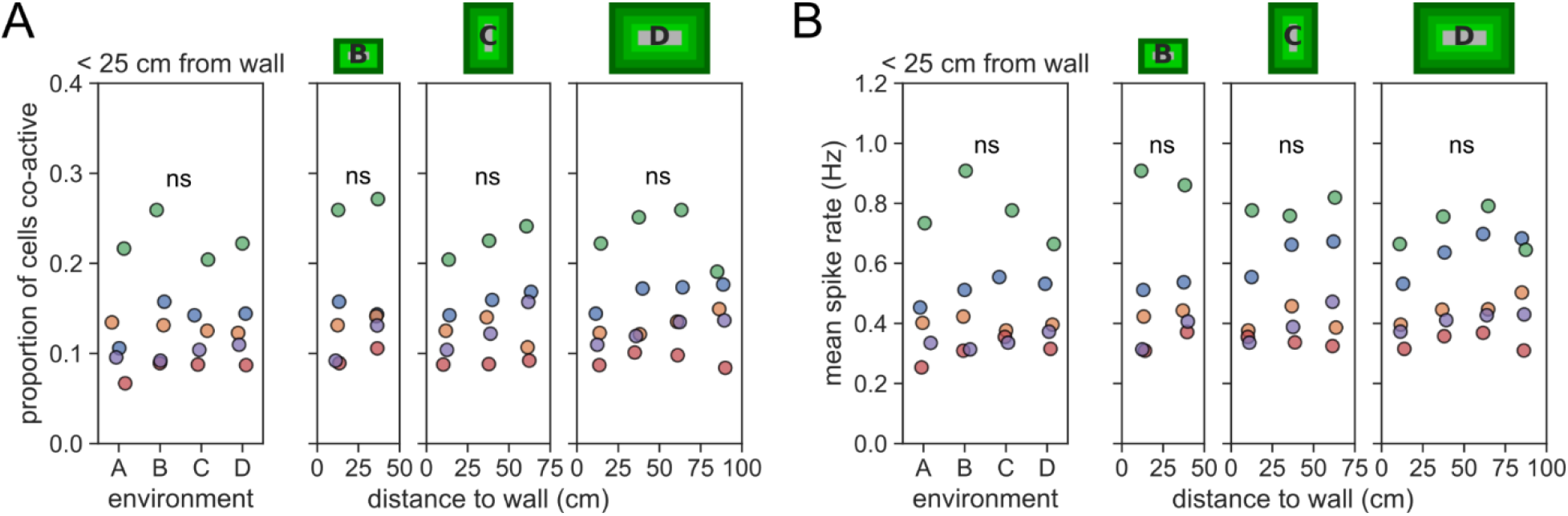
Place cell population statistics are constant within and between environments. (**A**) The proportion of place cells recorded in each animal that were firing at greater than 1 Hz was constant at different distances to the wall in all environments and same across environments. (**B**) The mean firing rate of all place cells recorded in each animal was constant at different distances to the wall in all environments and across environments.

**Fig. S10.**
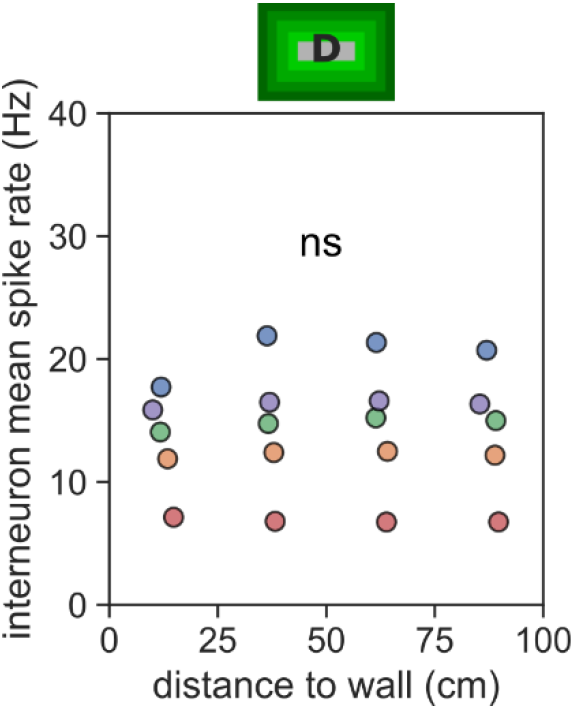
Interneuron firing rate is constant at all distances to walls. Mean spike rate of all interneurons detected in each animal at different distances to wall in environment D.

**Fig. S11.**
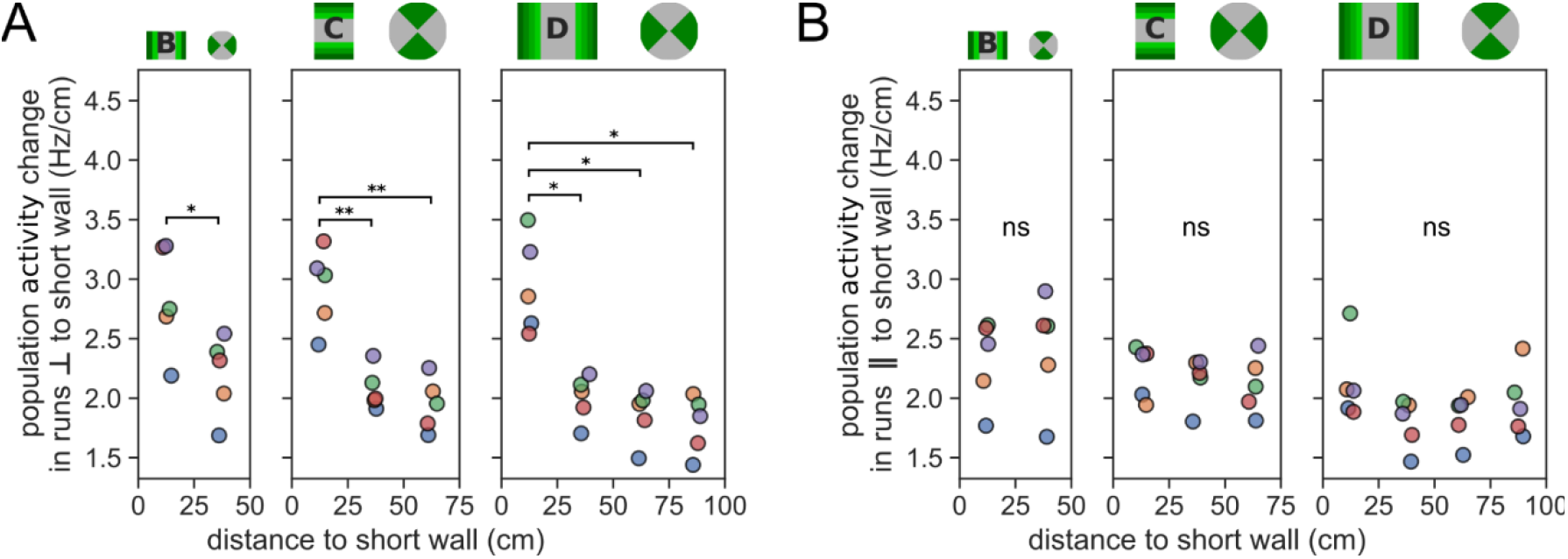
Population activity change was higher in runs orthogonal and close to walls in all environments. (**A**) Euclidean distance between vectors representing population activity at different distances to the short walls of each environment, only including runs orthogonal to the short wall. Pair-wise post hoc tests adjusted for multiple comparisons using Benjamini/Hochberg (non-negative) correction. (**B**) Same as **A**, but runs parallel to the short wall.

**Fig. S12.**
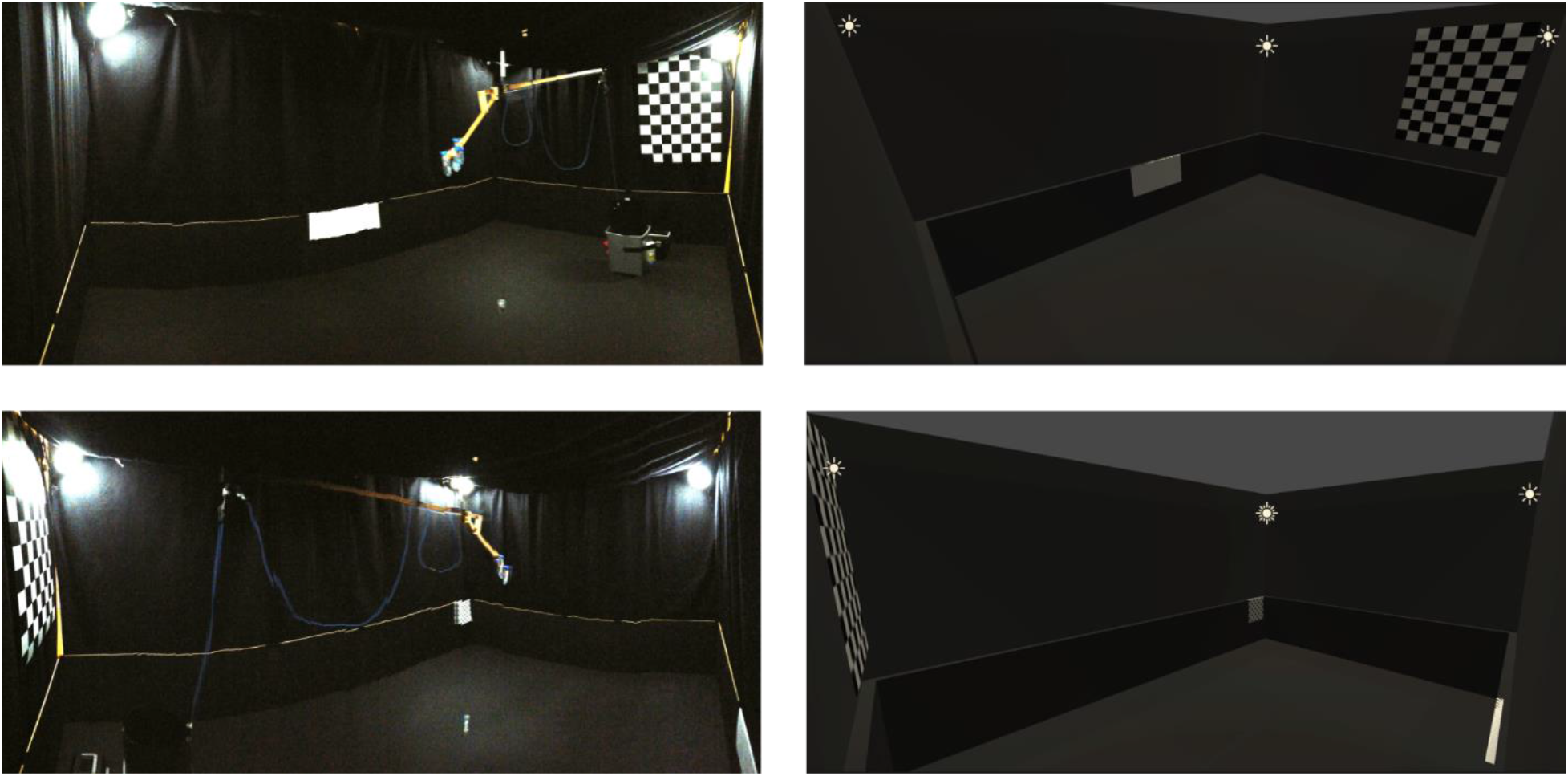
The virtual environment used to calculate visual change. Recording environment D (left) was replicated in a virtual environment (right) in order to estimate each rodent’s change in visual scene during its movement through the experimental environment.

